# mRNA-1273.262 and mRNA-1283.262 Vaccines Expressing XFG Antigens Neutralize Relevant Circulating Variants

**DOI:** 10.64898/2026.11.18.767652

**Authors:** Arshan Nasir, Clinton Ogega, Diana Wing Lee, Ziqiu Wang, Nathen Bopp, Emily King-Bursaw, Philippa Putnam, Sharanya Iyer, Lisa A Crawford, Amy Gorrie, Veronica Ritchie, Michael F Bender, Arvind Sharma, Tessa Speidel, Fadi Ghantous, Heather Davis, Laura Avena, Nicholas J Amato, Kai Wu, William R Scheif, Kiera Hohn, Carole Henry, Andrea Carfi, Rahnuma Wahid, Darin K Edwards, Alec W Freyn

## Abstract

SARS-CoV-2 continues to circulate and causes substantial disease burden as an endemic respiratory pathogen. Vaccination is a key public health measure to ameliorate the effects of COVID-19 disease but requires annual update to match strains that are circulating. Monitoring of circulating virus and characterization of matched candidate vaccines helps to inform which antigens will provide the greatest benefit to public well-being. The 2026-2027 season will leverage XFG as the variant of interest for vaccine production and *in vitro* and *in vivo* data was generated preclinically to support this selection. XFG variant vaccines express well in mRNA format as both full-length spike and linked subdomain antigen formats. Animals immunized with these vaccines elicit robust neutralizing responses to relevant JN.1-family virus variants in both primary series and booster models. In future seasons, robust epidemiologically monitoring including viral genome sequencing will improve early detection of relevant variants and subsequent preparation of vaccine antigens.

## Introduction

Severe acute respiratory syndrome coronavirus 2 (SARS-CoV-2) has caused more than 7 million reported deaths worldwide (WHO, June 2026).^1^ Beyond mortality, SARS-CoV-2 infection can result in hospitalization and downstream effects, with potentially long-lasting impact on the health and quality of life.^2^ Utilization of vaccines and other public health measures for the prevention of infection is critical for reducing the burden of COVID-19 disease.^3,4^ To optimize protection, SARS-CoV-2 vaccines must be periodically updated as the virus mutates and evades pre-existing population immunity.^5,6^ A preclinical pipeline for rapid identification and response to potentially immune-evasive emerging variants is therefore necessary to monitor, predict, and perform initial derisking on novel variants which are subject for use as vaccine constructs.

SARS-CoV-2 continues to evolve through a combination of antigenic drift and recombination, requiring periodic reassessment of vaccine antigen composition.^7,8^ Since the emergence of Omicron, successive vaccine updates have typically followed the dominant evolutionary trajectory of the virus, moving from ancestral and BA.4/BA.5-containing formulations (2020-22) to XBB.1.5 (2023-24), JN.1 and KP.2 (2024-25), and LP.8.1-based vaccines (2025-26). In 2025-26, LP.8.1 was selected as the vaccine antigen based on epidemiologic growth, antigenic risk assessment, and preclinical evidence that LP.8.1-matched mRNA vaccines broadened neutralization across LP.8.1 and other co-circulating JN.1-lineage viruses.^9–11^

The 2026 variant landscape presented a more complex epidemiologic pattern. Rather than a single lineage cleanly replacing its predecessors, multiple antigenically and phylogenetically distinct variants co-circulated, often in a region-specific manner. XFG, a recombinant derived from two JN.1-subvariants (LF.7 and LP.8.1.2), became dominant in North America and Europe in late summer and early Fall of 2025. XFG was designated a WHO Variant Under Monitoring in June 2025, as its global proportions increased.^12^ XFG remained the dominant variant in the United States in 2026, accounting for ∼68% of all genetic samples sequenced between April 12, 2026, and May 9, 2026.^13^ In parallel, another JN.1-derived variant NB.1.8.1 also increased in frequency, albeit its initial growth was mostly seen in Asian countries.^14^ As XFG wave slowed, NB.1.8.1 started to be detected more commonly outside Asia. . NB.1.8.1 has a more complicated evolutionary history, as it evolved from XDV, which was a recombinant between JN.1 and XBB viruses.^15^ The spike protein in NB.1.8.1 was inherited from JN.1. In parallel, BA.3.2, a highly divergent long-branch Omicron lineage with extensive spike divergence from both the ancestral and JN.1 lineage viruses, has been co-circulating alongside the JN.1-derived variants (XFG and NB.1.8.1) since at least March 2025, albeit at relatively lower levels but shows notable age-associated enrichment in pediatric populations.^16,17^ In addition to epidemiologic data and computational modeling, vaccine candidates were produced as mRNA-LNPs and characterized to ensure effective expression and immunogenicity. *In vitro* expression was assessed by mass spectrometry and flow cytometry, while immunogenicity was assessed *in vivo* in the murine model in the context of both pre-existing immunity and naïve animals. These data derisk the forward processing of vaccine candidates to a manufacturing-ready state and aid in understanding the breadth of immunogenicity elicited as well as the relevance of these responses to the contemporary variant landscape. With the WHO and major regulatory agencies allowing for the manufacture of XFG vaccine for the 2026/2027 COVID season, these preclinical data supported the ability of this vaccine to perform in the relevant markets for the target populations they are recommended.^19–21^

## Results

### Surveillance and structural analysis of SARS-CoV-2 variants

Routine genomic surveillance and risk assessment are performed to identify and characterize emerging SARS-CoV-2 variants with potential relevance for vaccine strain selection. Genomic and epidemiological data from SARS-CoV-2 infections worldwide are monitored throughout the year, integrating variant prevalence and growth dynamics with sequence-based and computational assessments of immune escape and other virologic features. Variants demonstrating concerning epidemiological growth, substantial antigenic change, or other features associated with increased risk are prioritized for further characterization.

Phylogenetic analysis illustrates the continued diversification of Omicron and the emergence of genetically distinct lineages, including contemporary JN.1- and BA.3.2-derived subclades (Figure 1A). Sequence-based assessment of variants of interest further characterizes amino acid changes at key antigenic sites, particularly within the receptor-binding domain (RBD) (Figure 1B). For example, XFG is a recombinant of LP.8.1.2 and LF.7, with a breakpoint in spike between amino acids 444 and 445, potentially combining immune-evasive mutations from both parental lineages.²²

**Figure 1.**
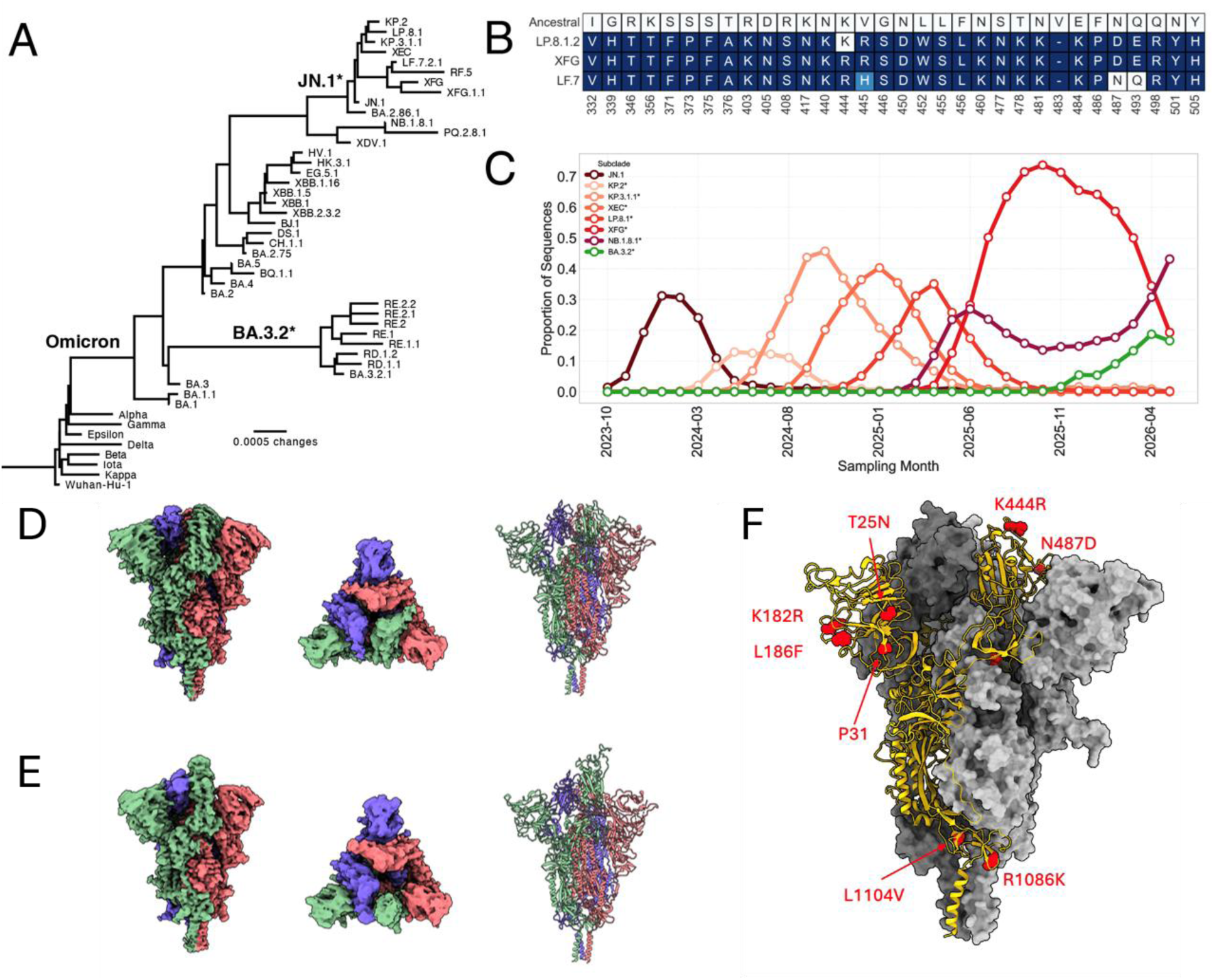
Evolutionary relationships, RBD diversity, and global circulation of contemporary SARS-CoV-2 variants. **(A)** Maximum-likelihood phylogeny of 47 representative SARS-CoV-2 genomes spanning the major variants and recombinant lineages that have emerged since the beginning of the COVID-19 pandemic. The tree was inferred from whole-genome sequences using the GTR+G4 substitution model and rooted on the branch leading to the ancestral Wuhan-Hu-1 strain. Major evolutionary transitions are labeled including Omicron and currently circulating JN.1 and BA.3.2* subclades. Tree topology and branch lengths are likely influenced by recombination. **(B)** RBD amino acid diversity among XFG, LP.8.1.2, LF.7, and ancestral RBDs. X-axis indicates amino acid numbering relative to the ancestral (Wuhan-Hu-1) strain. Mutations from the ancestral RBD are colored in different shades of blue. **(C)** Global monthly prevalence of major SARS-CoV-2 variants since late-2023. Variant prevalence was calculated as the proportion of sequences assigned to each variant group among all sequences collected during the corresponding calendar month. Asterisk indicates grouping of subvariants into the parental subclade, as assigned by nextclade. **(D and E)** Structural models of XFG full length spike with RBD down **(D)** or one RBD in the up conformation **(E).** From left to right, space-filling model, top-down view of space-filling model, ribbon diagram. **(F)** Full trimer space-filling model of the XFG spike with mutations from the LP.8.1 spike shown in red and highlighted on the model as red spheres. One monomer is shown as a ribbon diagram, with the gray space-filling models representing the other monomers.

These genetic changes occur against a backdrop of rapid turnover in the circulating variant landscape (Figure 1C). JN.1 predominated in early 2024 and was successively replaced by KP.2-, KP.3.1.1-, XEC-, and LP.8.1-derived variants. XFG subsequently expanded rapidly in mid-2025, reaching >70% global prevalence before declining in 2026. More recently, NB.1.8.1- and BA.3.2-derived viruses have increased in prevalence. Our computational risk assessments indicated that an XFG-based vaccine was likely to provide broad protection against the JN.1-lineage variants including (XFG and NB.1.8.1) and was also expected to be cross-reactive to BA.3.2 (Figure S1). 18 Integration of epidemiological trends with computational risk assessments is used to prioritize emerging variants for experimental evaluation and potential consideration as vaccine candidates. Based on its rapid global growth and distinct antigenic profile, XFG was prioritized for further characterization, including structural assessment of the spike protein by cryo-electron microscopy (**Figure 1D-F**). The spike bears several mutations relative to the prior season vaccine strain, LP.8.1, with key mutations in the RBD and N-terminal domain (NTD) likely contributing to immune escape.

### In vitro characterization of XFG vaccine candidates

To determine the ability of XFG vaccine candidates to express and generate similar amounts of protein antigen to previous strains, mass spectrometry and flow cytometry are utilized. Mass spectrometry of cells transfected with mRNA-LNP expressing either the full spike (1273) or RBD-NTD (1283) version of the XFG or LP.8.1 antigen were compared (**Figure 2A**). These data show similar peptide abundance of XFG when compared to LP.8.1 when the same version of the vaccine was used. Flow cytometry detects the amount of protein present on the cell surface at a fixed time after transfection with either full spike (**Figure 2B**) or RBD-NTD (**Figure 2C**) versions of the vaccine. Curves showing detection using either an S2-specific monoclonal antibody (mAb) or NTD-specific mAb were near overlapping for each variant when comparing XFG to LP.8.1.

**Figure 2.**
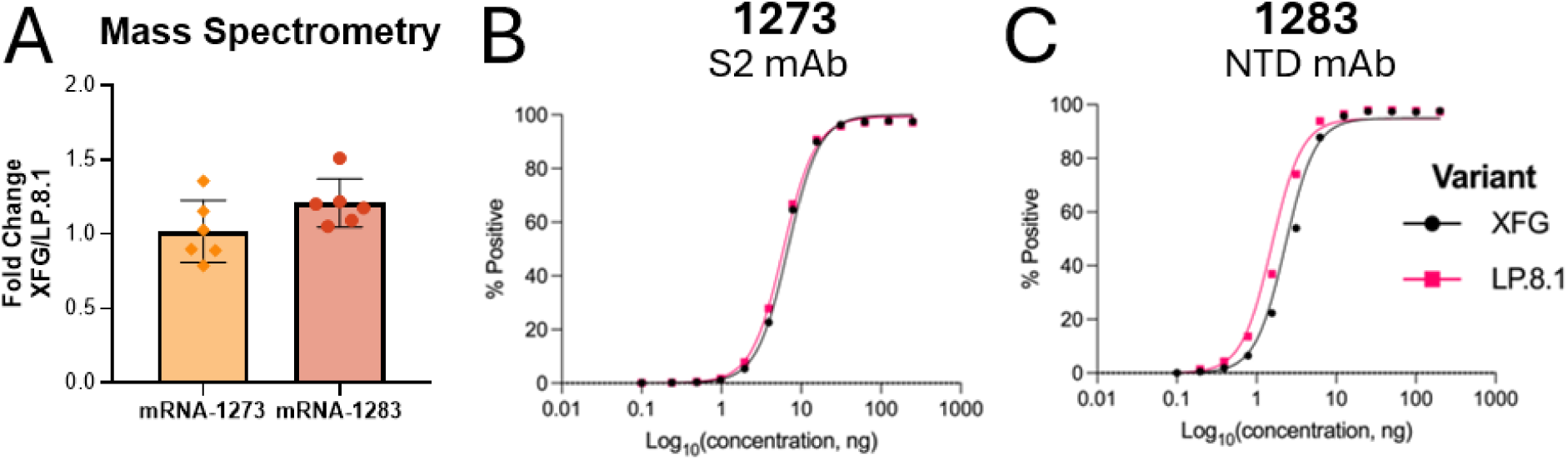
*In vitro* expression of new XFG candidate materials. **(A)** Mass spectrometry results of relevant peptides comparing protein abundance of XFG vaccine antigen relative to LP.8.1 vaccine antigen. Bar shows the average area under the curve of N=6 replicates with error showing standard deviation. **(B and C)** Flow cytometry of cells expressing vaccine antigen were assessed. Percent positive cells with detectable antigen expression are described for each dose of mRNA-LNP used. Detection reagents are listed in the title of each panel.

### In vivo immunogenicity of XFG vaccine candidates

Immunogenicity of vaccine candidates was assessed using murine models to assess the ability to generate binding and neutralizing titers in either naïve or pre-existing immunity conditions. Primary series studies were performed leveraging a two-dose regimen with immunization performed three weeks apart with homologous dosing (**Figure 3A**). Enzyme-linked immunosorbent assays (ELISAs) using ancestral Wuhan-Hu-1 full length spike protein were utilized to compare the amount of binding IgG antibodies elicited by the full-length spike and RBD-NTD vaccine designs (**Figure 3B and 3C**). Antibody titers elicited by the LP.8.1 vaccines were comparable (within 2-fold) to those generated by the XFG candidate vaccines. Neutralization titers were measured through the use of pseudoviruses expressing the SARS-CoV-2 spike from selected variants (**Figure 3D and 3E**). After primary series immunization, neutralization titers were comparable (within 2-fold) against the variants tested. Importantly, both vaccines elicited cross-neutralization responses against the distant BA.3.2.2 variant tested.

**Figure 3.**
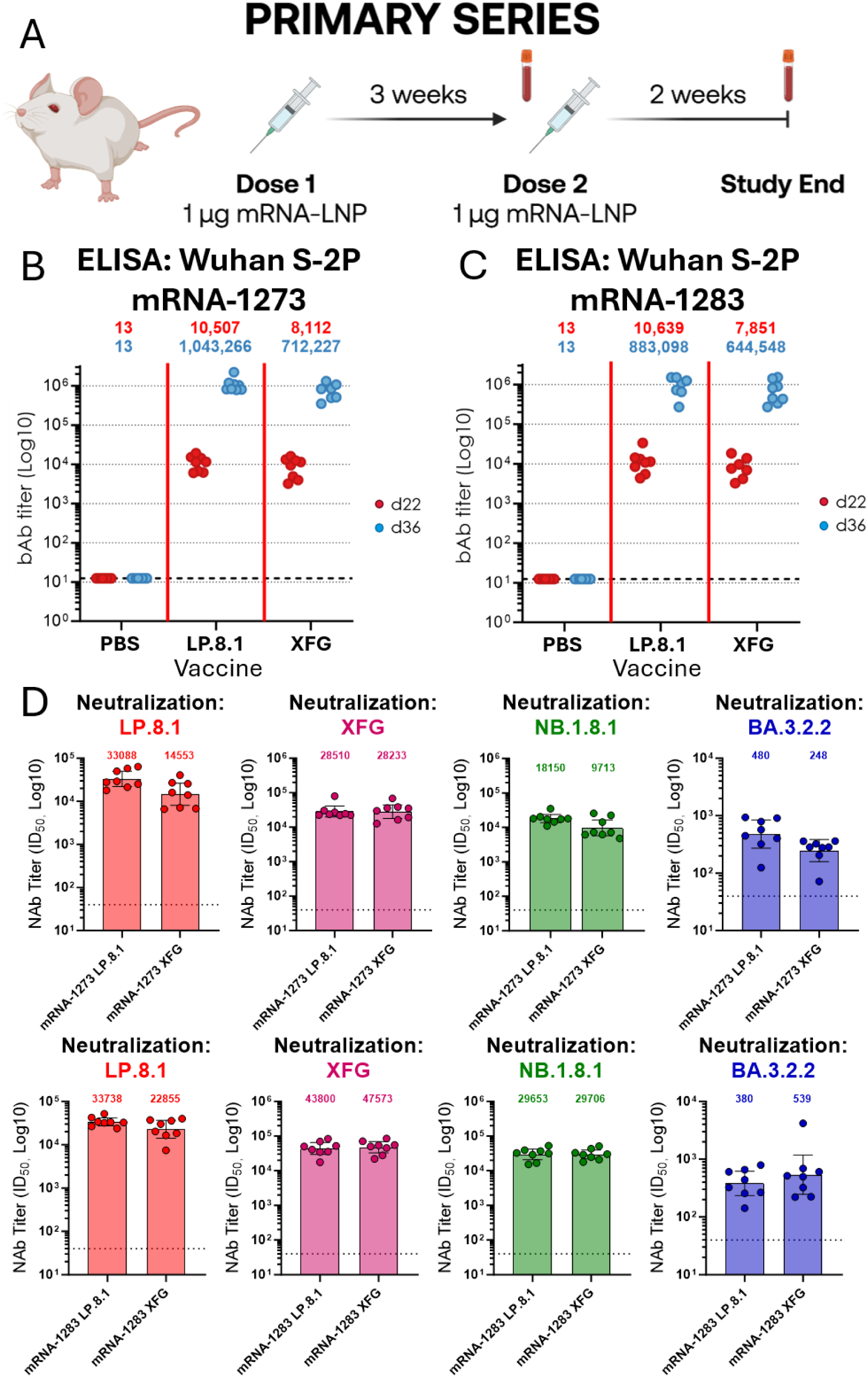
Primary series murine immunogenicity of XFG-based mRNA-LNP vaccine candidates. **(A)** Study schema highlighting dates of immunization and serum collection for primary series studies. **(B and C)** IgG binding titers are shown after primary series immunization with mRNA-1273 **(B)** or mRNA-1283 **(C)** based vaccines. Individual animal responses are shown for each timepoint with GMTs shown above each vaccine group. One half the LOD is shown as a dotted line on each graph. D22 – 22 days post prime, d36 – 36 days post prime. **(D and E)** Neutralization titers against VSV pseudoviruses expressing the spike of each variant listed on the x-axis are shown for each individual animal using day 36 sera. Bars represent GMT with error bars representing 95% confidence intervals. GMTs are also listed above each bar. LOD is shown as a dotted line.

Booster series studies were also leveraged to assess immunogenicity. These studies involved priming animals with a two-dose heterologous vaccine regimen comprised of a total of four vaccines representing the full-length spikes of major variant families from 2020-2025 in an attempt to model pre-existing immunity found in the general population (**Figure 4A**). After around two months, animals were then boosted with vaccine candidates based on the LP.8.1 or XFG variants. ELISAs were performed to assess binding antibody titers pre- and post-boost (**Figure 4B and 4C**). IgG binding titers were similar prior to boost (within 2-fold) though were found to be higher after booster immunization with either XFG vaccine than their cognate LP.8.1 vaccine (greater than 2-fold). Pseudovirus neutralization was also tested pre- and post-booster vaccination, and prior to boost all titers were similar (within 2-fold) when tested against the same pseudovirus (**Figure 4D and 4E**). XFG immunization was found to elicit higher neutralization titers post-boost than LP.8.1 vaccination against all pseudoviruses tested.

**Figure 4.**
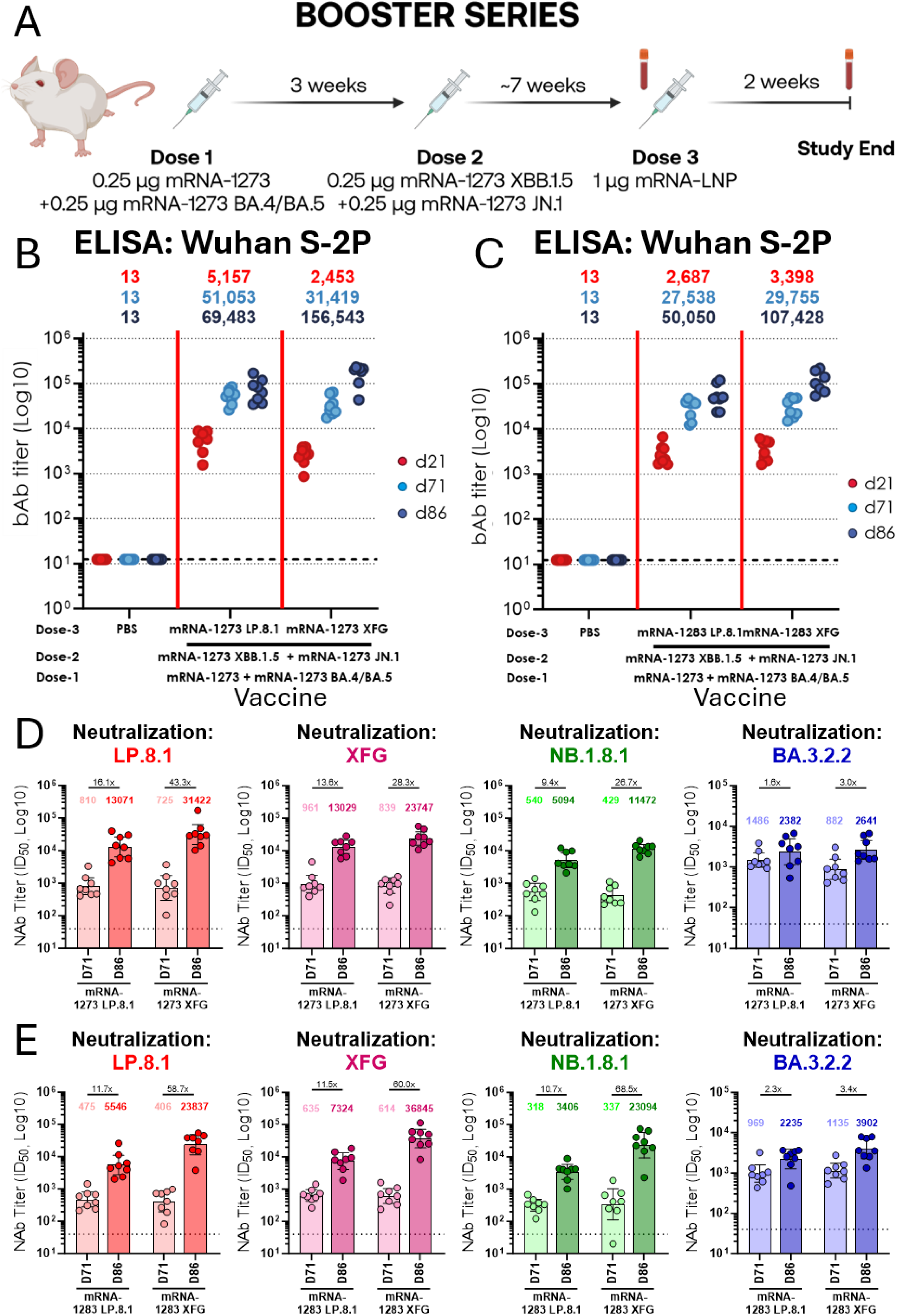
Booster series murine immunogenicity of XFG-based mRNA-LNP vaccine candidates. **(A)** Study schema highlighting dates of immunization and serum collection for booster series studies. **(B and C)** IgG binding titers are shown after booster series immunization with mRNA-1273 **(B)** or mRNA-1283 **(C)** based vaccines. Individual animal responses are shown for each timepoint with GMTs shown above each vaccine group. One half the LOD is shown as a dotted line on each graph. **(D and E)** Neutralization titers against VSV pseudoviruses expressing the spike of each variant listed on the x-axis are shown for each individual animal using pre-boost (day 71) and post-boost sera (day 86). Bars represent GMT with error bars representing 95% confidence intervals. GMTs and fold change are also listed above each bar. LOD is shown as a dotted line.

## Discussion

Surveillance of circulating SARS-CoV-2 variants and preclinical characterization of relevant vaccine candidates in both *in vitro* and *in vivo* settings are part of a broader continuum through clinical assessment and real-world evidence gathering that supports the use of seasonally strain-matched vaccines to confer optimal protection. Continued efforts to select the appropriate strain for the 2026/2027 respiratory season has led to the consensus that XFG would behave well as a vaccine antigen and provide relevant immune stimulation to drive antibody responses against strains predicted to circulate in the upcoming season. These preclinical findings will be complimented through clinical and real-world effectiveness data to ensure that the process to identify and characterize vaccine candidates is robust for future seasons.

XFG was predicted to be the candidate able to provide the greatest protection to the population. Assessment of this variant as both mRNA-1273 (full spike) and mRNA-1283 (RBD and NTD design) confirmed the ability of XFG candidate vaccines express well and demonstrated robust immunogenicity against the vaccine matched strain, allowing the consideration of these vaccines for commercial development. XFG antigens expressed in either vaccine format showed consistent trends for eliciting expression or immunogenicity data, providing confidence in this vaccine antigen as a candidate.

Additionally, both vaccines were shown to elicit neutralizing responses to JN.1-family variants and with cross-reactivity to BA.3.2.2. Though BA.3.2.2 exists as a long branch variant predicted to be antigenically distinct to the JN.1 family, the ability of XFG vaccines assessed to provide some level of cross-neutralization aided the selection of this variant as relevant to the upcoming season. The continued emergence of highly divergent strains, such as BA.3.2.2, serves as an example for the necessity of rapid testing and preclinical characterization of vaccine strains regardless of timing relative to season or strain selection.

Updating vaccine strains is necessary due to the consistent antigenic drift and recombination of SARS-CoV-2 viruses, potentially causing incremental loss of neutralization capability and immune recognition of novel variants. Further, long-branch variants such as BA.3.2.2 with substantial new mutations relative to currently circulating variants can constitute an antigenic shift where substantial pre-existing immunity is lost, leaving populations vulnerable to infection and the consequences inherent to each viral challenge.^23^ Continued surveillance and update of vaccine variants has shown an incremental increase in protection against contemporaneously circulating strains, suggesting that strain update, derisked by preclinical and clinical characterization, has a demonstrable impact on overall public health outcomes related to SARS-CoV-2 disease.

## Materials and Methods

### Phylogenetic analysis

A maximum-likelihood phylogeny was constructed using whole-genome sequences representing 47 major SARS-CoV-2 variants and recombinant lineages spanning the evolution of the virus from the start of the COVID-19 pandemic through June 2026. Representative genomes were manually selected to capture the major evolutionary transitions and dominant circulating variants. Sequences were aligned using Nextclade CLI (ver. 3.21.2, dataset: Nextstrain SARS-CoV-2 Wuhan-Hu-1 ORFs), and phylogenetic inference was performed with IQ-TREE 3 under the GTR+G4 nucleotide substitution model using automatic thread selection. The resulting tree was rooted using the Wuhan-Hu-1 reference genome and visualized to illustrate the evolutionary relationships among major SARS-CoV-2 lineages.

### Receptor-binding domain haplotypes

Consensus amino acid haplotypes of the spike RBD (residues 320–528, Wuhan-Hu-1 numbering) were generated for XFG, LP.8.1.2, and LF.7 Amino acid substitutions were determined relative to the ancestral Wuhan-Hu-1 reference spike RBD sequence (GenBank accession Id: MN908947) to illustrate the recombinant evolutionary history of XFG RBD.

### Variant prevalence analysis

SARS-CoV-2 metadata were downloaded from the GISAID EpiCoV database on 7 June 2026. Analyses were restricted to human-derived sequences with sample collection dates between November 1, 2023, and May 27, 2026. Metadata were processed using custom Python scripts to standardize collection dates and assign sequences to major variant groups based on Pango lineage designations and designated subclades, as defined by nextstrain. Variant groups included BA.3.2*, JN.1, KP.2*, KP.3.1.1*, LP.8.1*, NB.1.8.1*, XEC*, and XFG*. For each calendar month, variant prevalence was calculated as the number of sequences assigned to each variant group divided by the total number of sequences collected during that month. Monthly proportions were visualized using Python with the pandas and matplotlib libraries.

### Cryo-electron microscopy for structural resolution

Cryo-electron microscopy (cryo-EM) grids were prepared by applying 4 μL of purified XFG spike protein (0.5 mg/mL) to glow-discharged Quantifoil R1.2/1.3 300-mesh copper grids. Grids were blotted and vitrified by plunge-freezing into liquid ethane using a Vitrobot Mark IV (Thermo Fisher Scientific) operated at 4°C and 100% relative humidity. Cryo-EM data were collected on a Titan Krios G3i transmission electron microscope (Thermo Fisher Scientific) operating at 300 kV and equipped with a Gatan K3 direct electron detector and BioQuantum energy filter (Gatan) in zero-loss mode. Automated image acquisition was performed with EPU software at a nominal magnification of 105,000x, yielding a calibrated pixel size of 0.822 Å. Movies were acquired with a total electron dose of 51.5 e⁻/Å² distributed across 50 frames, using a nominal defocus range of −0.6 to −2.4 μm.

All movie stacks were processed in CryoSPARC v4.7.1. Raw movies underwent motion correction and contrast transfer function (CTF) estimation before automated particle picking with Topaz. Successive rounds of two-dimensional (2D) classification were used to remove non-particle images and retain high-quality particle projections. Three-dimensional (3D) classification was subsequently performed to evaluate conformational heterogeneity and eliminate remaining low-quality particles. The final reconstruction of the receptor-binding domain (RBD)-down and single RBD-up trimeric spikes were generated by non-uniform refinement with C3 symmetry using a dataset of 160,621 and 131,196 particles. Atomic model building was performed manually in Coot, followed by real-space refinement using the PHENIX.

### Flow cytometry cell-surface protein detection

Expi293F cells (Thermo Fisher Scientific) were used for all *in vitro* transfection studies. Cells were maintained in suspension culture in Expi293 Expression Medium (Thermo Fisher Scientific) at 37°C in a humidified incubator with 8% CO_2_ and orbital shaking at 125 rpm. For all experiments, cells were seeded at a density of 1 x 10⁶ cells/mL and verified to be in logarithmic growth phase with viability exceeding 97% at the time of transfection.

To evaluate *in vitro* protein expression, Expi293F cells were transfected with mRNA/LNP constructs encoding SARS-CoV-2 mRNA-1273 or mRNA-1283 LP.8.1 or XFG. A elevel-point, two-fold serial dilution beginning at 500 ng mRNA per 1 x 10⁶ cells was used for each construct. Following transfection, cells were incubated at 37°C with 8% CO₂ and shaking at 1,000 rpm and harvested approximately 24 hours later.

Cell surface expression of spike protein was assessed by flow cytometry using recombinant human ACE2 (hACE2), an NTD-specific monoclonal antibody (mAb) 118, and an S2-targeting mAb 103, all containing human Fc domains. Cells were stained with Alexa Fluor 647-conjugated goat anti-human IgG (Southern Biotech) together with Aqua viability dye (Thermo Fisher Scientific).

Flow cytometric data were acquired on an iQue3 instrument (Sartorius) using a sip rate of 1.5 μL/s for 5 seconds per well. Data acquisition was performed with ForeCyt software (Sartorius), and analyses were conducted using a sequential gating strategy of cells, singlets, viable cells, and antigen-positive cells. Protein expression was quantified as the percentage of positive cells. Graph generation and statistical analyses were performed using GraphPad Prism (GraphPad Software).

### Mass spectrometry for whole cell peptide detection

Samples were thawed and lysed using 5% SDS (Fisher Scientific, BP1311-1) prepared in MS-grade water (Fisher Scientific, W6-4) supplemented with protease inhibitors (Roche, 04 693 159 001), followed by sonication. Protein digestion was performed using a semi-automated workflow on the KingFisher Apex platform (Thermo Fisher Scientific) with MS-grade trypsin (Pierce, 90057). Following digestion, samples were acidified to a final concentration of 0.1% formic acid (Fisher Scientific, A117-50) and stored at 4°C until LC-MS analysis.

Acidified peptide samples were separated on a 15 cm x 75 μm PepMap C18 column (Thermo Fisher Scientific, ES75150PN) coupled to a Vanquish Neo UHPLC system. Peptides were eluted using the following solvent gradient (A→B): 0–2% over 2 minutes, 2–27% over 21 minutes, 27–45% over 3 minutes, and 45–95% over 2 minutes, followed by column washing and re-equilibration. Mass spectrometric data were acquired on an Exploris 480 mass spectrometer (Thermo Fisher Scientific) operating in data-independent acquisition (DIA) mode. Full MS (MS1) spectra were acquired at 60,000 resolution, followed by 32 sequential 8 m/z isolation windows for MS/MS (MS2) acquisition at 15,000 resolution. The normalized collision energy was set to 27%, and the MS2 automatic gain control (AGC) target was set to 1000%. Raw data were processed in Spectronaut (version 18.2) using a custom FASTA database consisting of the UniProt human reference proteome (UP000005640) supplemented with the SARS-CoV-2 mRNA-1273 LP.8.1 and XFG spike protein amino acid sequences.

### Murine study considerations

Animal studies were conducted using eight-week-old female BALB/c mice (Jackson Laboratories). Mice were immunized by intramuscular injection into a single hind limb with either 0.5 or 1 μg of mRNA-LNP formulated vaccine or phosphate-buffered saline (PBS) as a control, administered in a total volume of 50 μL. Blood samples were collected by submandibular bleeding during the study, and terminal blood collection was performed by cardiac puncture under isoflurane anesthesia. A total of two cohorts, consisting of 40 animals, were included in these studies.

All animal procedures were conducted in accordance with Protocol 23-07-016, as approved by Moderna’s Institutional Animal Care and Use Committee (IACUC). All activities complied with the National Institutes of Health (NIH) guidelines, the Animal Welfare Act, and all applicable U.S. federal regulations. Euthanasia was performed by isoflurane inhalation followed by exsanguination under anesthesia and cervical dislocation, in accordance with the American Veterinary Medical Association (AVMA) Guidelines for the Euthanasia of Animals (2013 Report of the AVMA Panel on Euthanasia).

### Enzyme-linked immunosorbent assays for binding antibody detection

Binding antibody responses were measured by ELISA using 384-well microtiter plates (Thermo Fisher Scientific; Cat. # 4644718) coated with 50 μL of recombinant Wuhan-Hu-1 spike protein (S-2P). Plates were incubated overnight at 4°C and subsequently blocked with SuperBlock (Thermo Fisher Scientific; Cat. # 37536) for 1 hour at 37°C. Serum samples were serially diluted in 5% BSA in TBS (TBS5; Boston Bioproducts, IBB-187), added to the coated plates, and incubated for 1 hour at 37°C. Plates were then washed four times with PBST.

Goat anti-mouse IgG-HRP (Southern Biotech; Cat. #1030-05), diluted in TBS5, was added to each well, followed by incubation for 1 hour at 37°C. After four additional washes with PBST, TMB substrate (Thermo Fisher Scientific; Cat. # 34029) was added to develop the reaction, which was subsequently terminated by the addition of TMB stop solution (Invitrogen; Cat. #374489-000). Absorbance was measured at 450 nm, and antibody titers were calculated by fitting the data to a four-parameter logistic regression model in Genedata Screener (Genedata Inc.). Endpoint titers were defined as the reciprocal serum dilution corresponding to an optical density of approximately 1.0 at 450 nm, normalized to a mouse standard included on each plate.

### Vesicular stomatitis virus-based SARS-CoV-2 neutralization assays

Codon-optimized, full-length SARS-CoV-2 spike genes representing LP.8.1, XFG, or BA.3.2.2 were cloned into the pCAGGS expression vector. VSVΔG-based SARS-CoV-2 pseudoviruses were generated by transfecting BHK-21/WI-2 cells with the respective spike expression plasmids, followed by infection with VSVΔG-firefly-luciferase, as previously described.^24^ Neutralization assays were performed using A549 cells (ATCC; CCL-185) engineered to express human ACE2 and TMPRSS2 as target cells. These cells were maintained in DMEM supplemented with 10% fetal bovine serum (FBS). Prior to the assay, serum samples were heat-inactivated at 56°C for 45 minutes and serially diluted in DMEM containing 10% FBS. Diluted serum samples, or culture medium alone as the virus-only control, were incubated with VSVΔG-based SARS-CoV-2 pseudovirus at 37°C for 45 minutes. The resulting virus-serum mixtures were then added to A549-hACE2-TMPRSS2 cells and incubated for 18 hours at 37°C.

Following infection, One-Glo reagent (Promega; E6120) was added to the culture medium at a 1:1 volume ratio, and luminescence was measured using a PHERAstar FSX plate reader (BMG). Neutralization was calculated as the percentage reduction in relative light units compared with the virus-only control. Neutralization curves and titers were determined by fitting the data with a four-parameter logistic regression model in Prism 10 (GraphPad).

## Acknowledgements

Cryo-EM data was collected at the Cryo-EM Facility at MIT.nano. We acknowledge beamline 19-ID (NYX) at NSLS-II, a U.S. DOE Office of Science User Facility at Brookhaven National Laboratory (Contract DE-SC0012704) and thank Kevin Battaile for X-ray data collection. We thank Dr. Michael Whitt and Rita Kansal for support on pseudovirus production. We gratefully acknowledge all data contributors, i.e., the Authors and their Originating laboratories responsible for obtaining the specimens, and their Submitting laboratories for generating the genetic sequence and metadata and sharing via the GISAID Initiative, on which some of this research is based. All funding was provided by Moderna Inc.

## References

1. WHO. WHO COVID-19 dashboard. May 9, 2026. Accessed June 3, 2026. https://data.who.int/dashboards/covid19/deaths

2. Hou Y, Gu T, Ni Z, Shi X, Ranney ML, Mukherjee B. Global Prevalence of Long COVID, Its Subtypes, and Risk Factors: An Updated Systematic Review and Meta-analysis. Open Forum Infect Dis. 2025;12(9):ofaf533. doi:10.1093/ofid/ofaf533

3. Scott J, Abers MS, Marwah HK, et al. Updated Evidence for Covid-19, RSV, and Influenza Vaccines for 2025–2026. N Engl J Med. 2025;393(22):2221–2242. doi:10.1056/nejmsa2514268

4. Liu Y, Diamond C, Abbott S, et al. The Impact of Public Health and Social Measures (PHSMs) on SARS-CoV-2 Transmission in the WHO European Region (2020–2022). Influ Other Respir Viruses. 2024;18(12):e70036. doi:10.1111/irv.70036

5. Link-Gelles R, Rowley EAK, Irving SA, et al. Estimated 2023-2024 COVID-19 Vaccine Effectiveness in Adults. JAMA Netw Open. 2025;8(6):e2517402. doi:10.1001/jamanetworkopen.2025.17402

6. Levy ME, Chilunda V, Heaton PR, et al. XBB.1.5 mRNA COVID-19 vaccine protection against inpatient or emergency department visits among adults infected with SARS-CoV-2 JN.1 and XBB-lineage variants. Front Immunol. 2025;16:1470609. doi:10.3389/fimmu.2025.1470609

7. Bolze A, Basler T, White S, et al. Evidence for SARS-CoV-2 Delta and Omicron co-infections and recombination. Med. 2022;3(12):848–859.e4. doi:10.1016/j.medj.2022.10.002

8. Yewdell JW. Antigenic drift: Understanding COVID-19. Immunity. 2021;54(12):2681–2687. doi:10.1016/j.immuni.2021.11.016

9. FDA. COVID-19 Vaccines (2025-2026 Formula) for Use in the United States Beginning in Fall 2025. August 12, 2025. Accessed May 22, 2025. https://www.fda.gov/vaccines-blood-biologics/industry-biologics/covid-19-vaccines-2025-2026-formula-use-united-states-beginning-fall-2025

10. EMA. EMA recommendation to update the antigenic composition of authorised COVID-19 vaccines for 2025-2026. May 17, 2025. Accessed August 12, 2025. https://www.ema.europa.eu/en/documents/other/ema-recommendation-update-antigenic-composition-authorised-covid-19-vaccines-2025-2026_en.pdf

11. WHO. Statement on the antigen composition of COVID-19 vaccines. May 15, 2025. Accessed August 12, 2025. https://www.who.int/news/item/15-05-2025-statement-on-the-antigen-composition-of-covid-19-vaccines

12. WHO. WHO TAG-VE Risk Evaluation for SARS-CoV-2 Variant Under Monitoring: XFG. June 25, 2025. Accessed August 24, 2026. https://www.who.int/publications/m/item/risk-evaluation-for-sars-cov-2-variant-under-monitoring-xfg

13. CDC. Variants and Genomic Surveillance. August 28, 2025. Accessed June 29, 2026. https://www.cdc.gov/covid/php/variants/variants-and-genomic-surveillance.html

14. Pranto AH, Promi AT, Islam S, Islam MdR. Resurgent COVID-19 Wave in India and South Asia: Is the NB.1.8.1 Variant a Renewed Global Health Concern? Heal Sci Rep. 2025;8(11):e71453. doi:10.1002/hsr2.71453

15. Uriu K, Okumura K, Uwamino Y, et al. Virological characteristics of the SARS-CoV-2 NB.1.8.1 variant. Lancet Infect Dis. 2025;25(8):e443. doi:10.1016/s1473-3099(25)00356-1

16. Jule Z, Römer C, Hossen T, et al. Evolution and viral properties of the SARS-CoV-2 BA.3.2 subvariant. Virus Evol. 2026;12(1):veag011. doi:10.1093/ve/veag011

17. Wu M, Hong H, Guo Y, et al. SARS-CoV-2 BA.3.2.2 is more evasive of neutralisation by plasma from young children. Lancet Infect Dis. 2026;26(9):e328–e329. doi:10.1016/s1473-3099(26)00349-x

18. Nasir A, Lee D, Avena LE, et al. Predictive modeling of immune escape and antigenic grouping of SARS-CoV-2 variants. J Virol. Published online 2026:e0022526. doi:10.1128/jvi.00225-26

19. EMA. ETF recommends updating COVID-19 vaccines to target XFG variant. May 29, 2026. Accessed August 24, 2026. https://www.ema.europa.eu/en/news/etf-recommends-updating-covid-19-vaccines-target-xfg-variant

20. WHO. Statement on the antigen composition of COVID-19 vaccines. May 16, 2026. Accessed August 24, 2026. https://www.who.int/news/item/16-05-2026-statement-on-the-antigen-composition-of-covid-19-vaccines

21. FDA. COVID-19 Vaccines (2026-2027 Formula) for Use in the United States Beginning in Fall 2026. May 29, 2026. Accessed August 24, 2026. https://www.fda.gov/vaccines-blood-biologics/industry-biologics/covid-19-vaccines-2026-2027-formula-use-united-states-beginning-fall-2026

22. Guo C, Yu Y, Liu J, et al. Antigenic and virological characteristics of SARS-CoV-2 variants BA.3.2, XFG, and NB.1.8.1. Lancet Infect Dis. 2025;25(7):e374–e377. doi:10.1016/s1473-3099(25)00308-1

23. Carabelli AM, Peacock TP, Thorne LG, et al. SARS-CoV-2 variant biology: immune escape, transmission and fitness. Nat Rev Microbiol. 2023;21(3):162–177. doi:10.1038/s41579-022-00841-7

24. Whitt MA. Generation of VSV pseudotypes using recombinant ΔG-VSV for studies on virus entry, identification of entry inhibitors, and immune responses to vaccines. J Virol Methods. 2010;169(2):365–374. doi:10.1016/j.jviromet.2010.08.006

