## Supplemental Figures for "mRNA-1273.262 and mRNA-1283.262 Vaccines Expressing XFG Antigens Neutralize Relevant Circulating Variants"

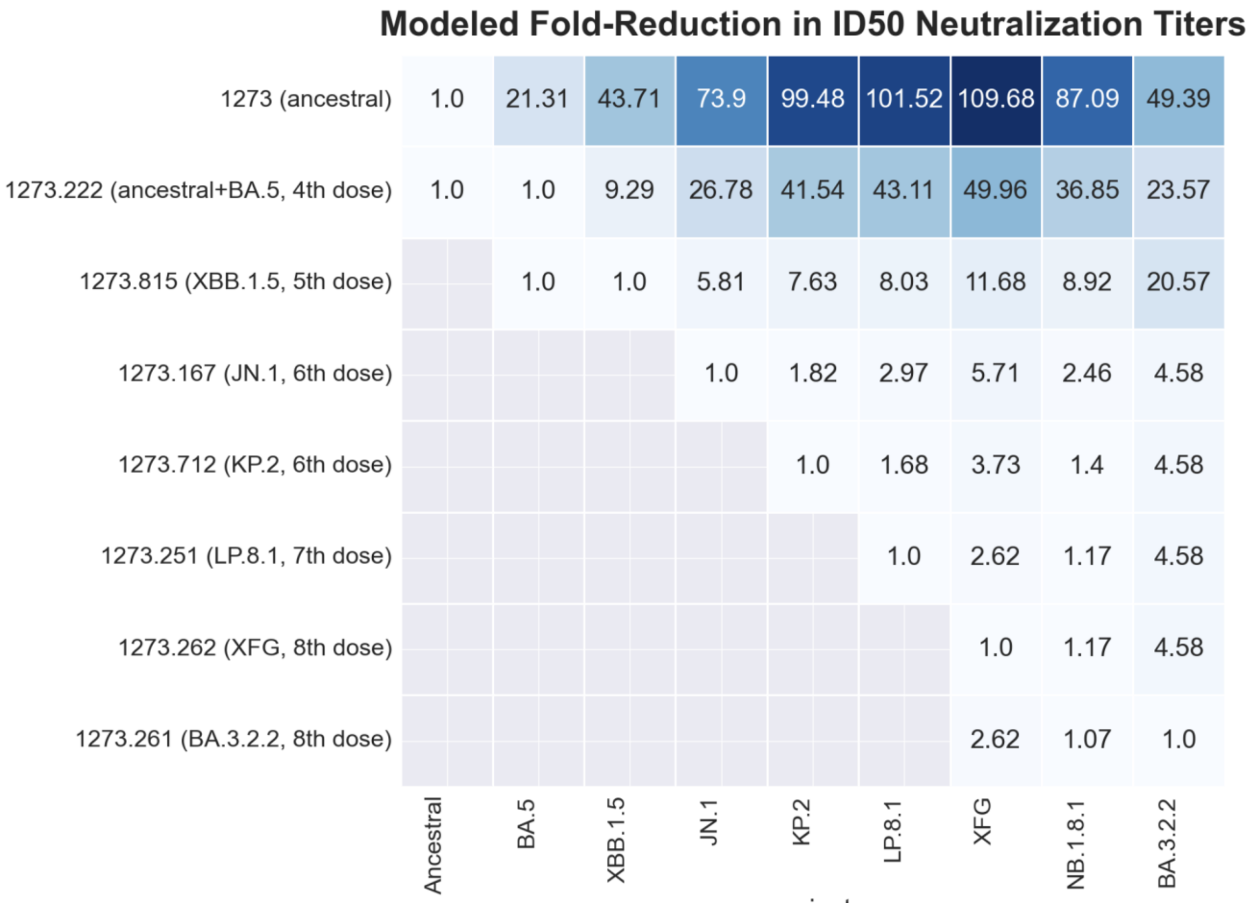


**Figure S1.** The matrix lists the predicted immune escape for variants against different immunization backgrounds. The values inside the matrix are the predicted fold-drop in theID50 neutralizing antibody GMTs on the vaccine strain, derived from the statistical model.[18]
